# Striatal dopamine and serotonin release operate largely in parallel with limited, asymmetric crosstalk

**DOI:** 10.1101/2025.03.28.645922

**Authors:** Chaudun Fabrice, Yu Liu, Juan Enriquez Traba, Agnès Hiver, Christian Lüscher

## Abstract

Dopamine and serotonin jointly regulate striatal functions that support reinforcement, decision-making and motivated behavior, yet whether activity in one system directly recruits release from the other remains unclear. Using anatomical tracing, cell type-specific optogenetics and genetically encoded fluorescent sensors, we measured dopamine and serotonin dynamics in the nucleus accumbens and dorsal striatum after cocaine administration and selective neuronal stimulation. Activation of ventral tegmental area dopamine neurons increased dopamine release without affecting serotonin. In contrast, activation of dorsal raphe serotonergic neurons robustly increased serotonin and weakly entrained dopamine signals without producing a sustained increase in dopamine amplitude. These findings indicate that striatal dopamine and serotonin release operate largely in parallel, with limited and directionally asymmetric crosstalk.

## Main Text

Midbrain dopamine (DA) neurons innervate the striatum from the ventral tegmental area (VTA) and substantia nigra, whereas tryptophan hydroxylase-positive neurons in the dorsal raphe nucleus (DRN) provide a major source of striatal serotonin (5-HT) (1,2). Dysregulation of either system contributes to neuropsychiatric and neurodegenerative disorders, including depression, schizophrenia, addiction and Parkinson’s disease (3–5). Despite their overlapping projections and behavioral functions, we still do not know how these neuromodulators influence one another in the striatum.

One influential model proposes that DA and 5-HT act as opponent systems: DA promotes behavioral activation and reinforcement, whereas 5-HT restrains responding or signals unfavorable outcomes (6–8). Pharmacological manipulations support this view by producing contrasting effects on reward processing, feeding and intracranial self-stimulation (9–12). Recent fiber-photometry recordings also revealed inverse striatal DA and 5-HT dynamics during reinforcement (13) and the transition from controlled to compulsive cocaine use (14). Moreover, 5-HT can inhibit DA neurons in the VTA and substantia nigra and reduce striatal DA release (15,16).

Other findings instead indicate cooperation. Local 5-HT infusion increases extracellular striatal DA (17), and serotonergic manipulations can enhance DA-dependent reinforcement. Selective 5-HT reuptake inhibitors alter brain-stimulation reward thresholds and potentiate cocaine reinforcement, whereas 5-HT depletion weakens cocaine reward and disrupts the effects of amphetamine on impulsivity (18–21). DRN activity may also encode reward information complementary to that carried by DA neurons (22).

Together, these conflicting observations suggest that DA–5-HT interactions depend on circuit context and experimental timescale, but they do not establish whether activity in one neuromodulatory system directly recruits transmitter release from the other.

Previous studies largely relied on pharmacology, lesions, electrical stimulation or slow neurochemical measurements (23). These approaches could not selectively activate one neuromodulatory population while resolving rapid release of both transmitters in defined striatal regions. This limitation leaves a fundamental question unanswered: does activity in one system directly recruit transmitter release from the other, or do DA and 5-HT operate mainly through parallel circuits?

Here, we combined projection mapping, cell type-specific optogenetics and genetically encoded fluorescent sensors to measure DA and 5-HT dynamics in the nucleus accumbens (NAc) and dorsal striatum (DS) of freely moving mice. Selective activation of VTA DA neurons increased striatal DA without altering 5-HT. In contrast, activation of DRN serotonergic neurons strongly increased 5-HT and imposed only a small, temporally synchronized DA component. These results reveal a directionally asymmetric interaction in which striatal 5-HT operates independently of VTA DA activity, whereas DRN serotonergic activity exerts a limited influence on striatal DA signaling.

## Results

### Cocaine evokes distinct dopamine and serotonin dynamics across the striatum

We first measured cocaine-evoked DA and 5-HT signals in the NAc and DS of freely moving mice in an open field arena. We expressed dLight1.2 or 5-HT3.0 in both regions and recorded fluorescence before and after an intraperitoneal injection (i.p.) of cocaine (15 mg kg⁻¹; Figure 1A–C).

**Figure 1:**
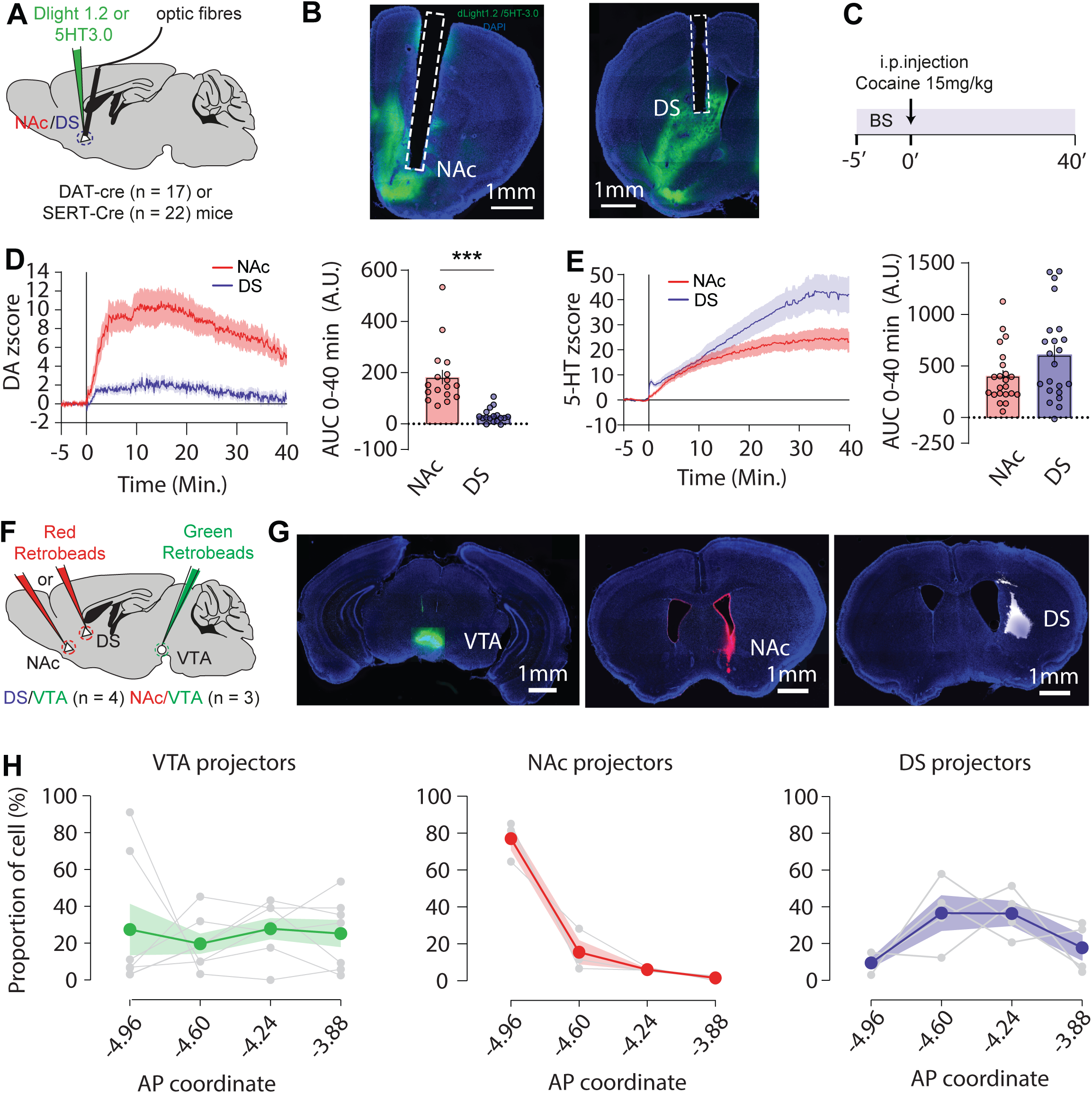
Temporal dynamics of DA and 5-HT release evoked by cocaine and topographical organization of DR projectors. **(A)** Schematics of mouse preparation to record DA or 5-HT release in NAc and DS region. **(B)** Representative histological example of viral expression and fibre implantation in NAc and DS. (**C**) Schematic of recording timeline following i.p. injection of cocaine 15mg/kg. **(D)** Left: z-score group data (Mean ± SEM) of NAc (red) and DS (Blue) DA release evoked by cocaine. Right: Quantification of AUC after i.p. injection of cocaine (Two-sided Mann-Whitney test, \*\*\**P*<0.001). **(E)** Left: z-score group data (Mean ± SEM) of NAc (red) and DS (Blue) 5-HT release evoked by cocaine. Right: Quantification of AUC after i.p. injection of cocaine (Two-sided Mann-Whitney test, *P=* 0.1310). **(F)** Schematic of retrobeads injection site used to map the topographical organization of DR projectors. **(G)** Representative examples of retrobeads injection site within the VTA, NAC and DS. **(H)** Proportion of cell (%) from VTA projectors (green), NAc projectors (red) and DS projectors on 4 different coronal planes.

Cocaine rapidly increased DA in the NAc, producing a large response that gradually declined during the recording period. Cocaine also increased DA in the DS, but the response was substantially smaller than in the NAc (Figure 1D). By contrast, cocaine progressively increased 5-HT in both regions. These signals remained elevated throughout the recording period, and their integrated amplitudes did not differ significantly between the NAc and DS (Figure 1E). Thus, cocaine produced regionally distinct DA responses but broadly similar and more sustained 5-HT responses.

We next mapped dorsal raphe (DR) neurons projecting to the NAc, DS and VTA using retrograde tracers (Figure 1F,G). NAc-projecting neurons concentrated in posterior DR sections, whereas DS-projecting neurons occurred mainly at intermediate rostrocaudal levels. VTA-projecting neurons showed a broader distribution across the DR (Figure 1H). These results reveal a topographical organization of DR outputs to striatal and midbrain targets.

### Activation of VTA dopamine neurons releases dopamine but not serotonin in the striatum

We next tested whether VTA DA-neuron activity recruits striatal 5-HT release. We expressed ChrimsonR in VTA DA neurons of DAT-Cre mice and stimulated these neurons with 2-Hz trains of 20-Hz bursts for 15 s while recording DA or 5-HT signals in the NAc and DS (Figure 2A,B).

**Figure 2:**
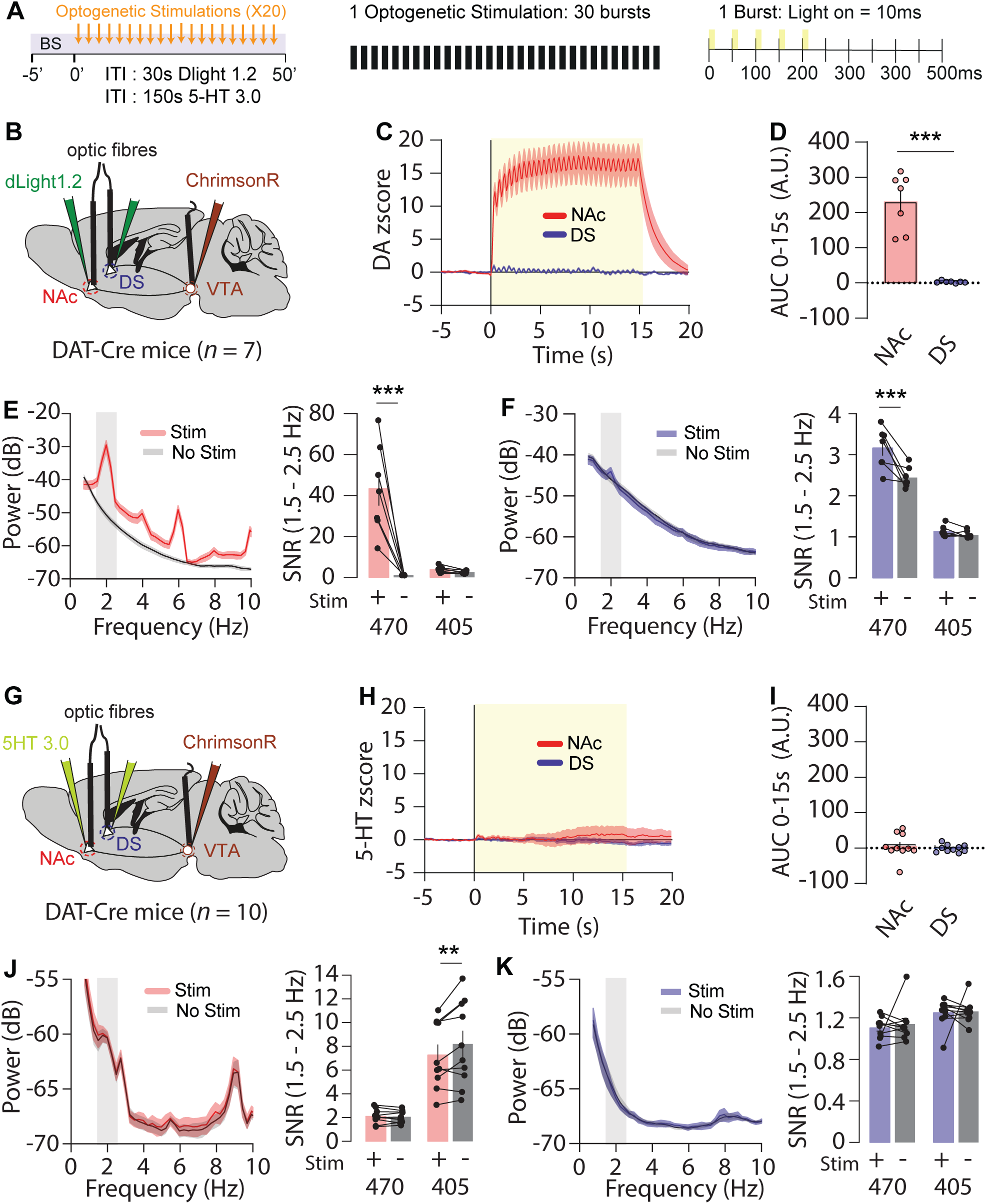
Stimulation of VTA DA neurons increases striatal DA but not 5-HT signals. **(A)** Left: Protocol of optogenetic stimulation used to assess DA and 5-HT release. Middle: Parameters of optogenetic stimulation consisted of 30 bursts at 2 Hz. Right: Each burst comprised five light stimulations of 10 ms each at 20 Hz every 500 ms. **(B)** Schematic of viral strategies used in DAT-Cre animals (*n* = 7) to simultaneously recorded DA release during optogenetic manipulation of VTA DA neurons. **(C)** z-score group data (Mean ± SEM) of NAc (red) and DS (blue) DA release evoked by optogenetic stimulation of VTA DA neurons. **(D)** Quantification of AUC during optogenetic stimulation (Two-sided Mann-Whitney test, \*\*\**P*<0.001). **(E, F)** Left: Group average power spectral density plot (PSD, Mean ± SEM) in decibel (dB) of DA ΔF/F signal during stimulation (red for NAc and blue for DS) versus non-stimulation epochs (grey). Right: 2 Hz signal-to-noise ratio quantification (SNR) of biosensor (470) and isosbestic (405) signal during stimulation versus non stimulation period (NAc : Two-way repeated measures ANOVA: Channel effect, F(_1,12_) = 21.02, ***P<0.001; Stimulation effect, F(_1,12_) = 27.95, ***P<0.001; Channel vs Stimulation, F(_1,12_) = 24.64, ***P<0.001; Bonferroni post hoc analysis, \*\*\**P*<0.001; DS: Two-way repeated measures ANOVA: Channel effect, F(_1,12_) = 191.7, ***P<0.001; Stimulation effect, F(_1,12_) = 19.21, ***P<0.001; Channel vs Stimulation, F(_1,12_) = 12.23, **P<0.01; Bonferroni post hoc analysis, \*\*\**P*<0.001). **(G)** Schematic of viral strategies used in DAT-Cre animals (*n* = 10) to simultaneously recorded 5-HT release during optogenetic manipulation of VTA DA neurons. **(H)** z-score group data (Mean ± SEM) of NAc (red) and DS (Blue) 5-HT release evoked by optogenetic stimulation of VTA DA neurons. **(I)** Quantification of AUC during optogenetic stimulation (Two-sided Mann-Whitney test, *P>*0.05) **(J,K)** Left: Group average power spectral density plot (PSD, Mean ± SEM) in decibel (dB) of 5-HT ΔF/F signal during stimulation (red for NAc and blue for DS) versus non-stimulation epochs (grey). Right: 2 Hz signal-to-noise ratio quantification (SNR) of biosensor (470) and isosbestic (405) signal during stimulation versus non stimulation period (NAc: Two-way repeated measures ANOVA: Channel effect, F(_1,18_) = 31.57, ***P<0.001; Stimulation effect, F(_1,18_) = 6.88, **P<0.01; Channel vs Stimulation, F(_1,18_) = 8.231, **P<0.01; Bonferroni post hoc analysis, \*\**P*<0.01; DS: Two-way repeated measures ANOVA: Channel effect, F(_1,18_) = 14.43, **P<0.01; Stimulation effect, F(_1,18_) = 0.25, P>0.05; Channel vs Stimulation, F(_1,18_) = 0.050, P>0.05).

VTA DA-neuron stimulation produced a rapid and sustained DA increase in the NAc but little change in the mean DA signal in the DS (Figure 2C,D). Spectral analysis revealed increased power restricted to the biosensor channel at the 2-Hz stimulation frequency in both regions, although the response was substantially larger in the NAc (Figure 2E,F). Thus, VTA stimulation strongly increased accumbal DA and weakly entrained DA signals in the DS.

The same stimulation did not alter mean 5-HT signals in either region (Figure 2G–I). It also failed to increase 5-HT signal power of the biosensor channel at 2 Hz in the NAc or DS (Figure 2J,K). VTA DA-neuron activation therefore drove striatal DA release without recruiting detectable 5-HT release.

### Activation of dorsal raphe serotonin neurons releases serotonin and weakly entrains dopamine signals

We then tested whether DR serotonergic activity influences striatal DA release. We expressed ChrimsonR in DR serotonergic neurons of SERT-Cre mice and applied the same stimulation protocol while recording 5-HT or DA signals in the NAc and DS (Figure 3A,B).

**Figure 3.**
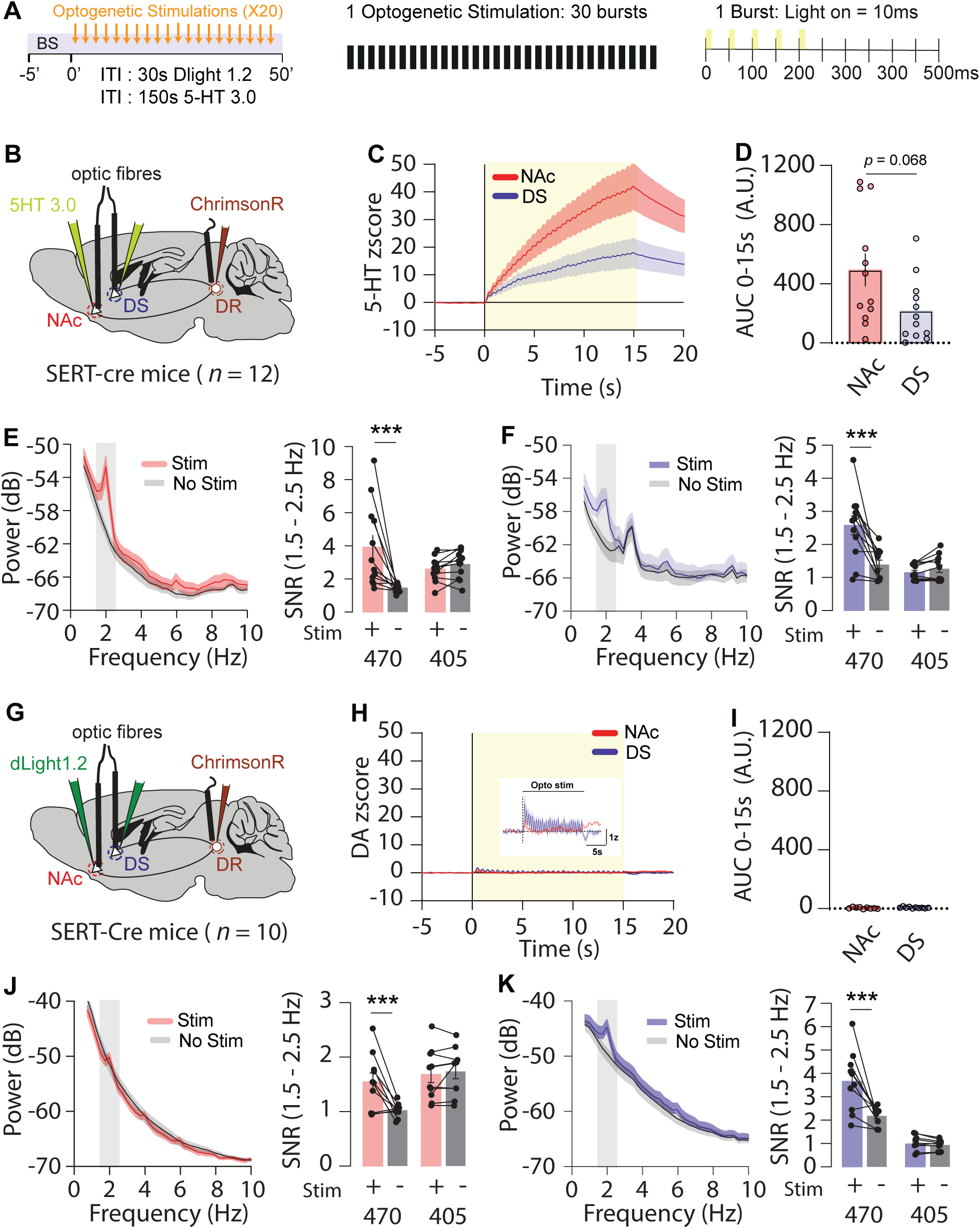
Stimulation of DRN 5-HT system increases 5-HT and synchronized DA signals in the striatum. **(A)** Left: Protocol of optogenetic stimulation used to assess DA and 5-HT release. Middle: Parameters of optogenetic stimulation realized consisting of 30 bursts at 2 Hz. Right: Each burst comprised of 5 light stimulations of 10 ms each at 20 Hz every 500 ms. **(B)** Schematic of viral strategies used in SERT-Cre animals (*n* = 12) to simultaneously recorded 5-HT release during optogenetic manipulation of DRN serotonergic neurons. **(C)** z-score group data (Mean ± SEM) of NAc (red) and DS (Blue) 5-HT release evoked by optogenetic stimulation of DRN serotonergic neurons. **(D)** Quantification of AUC during optogenetic stimulation (Two-sided Mann-Whitney test, *P* = 0.0684). **(E, F)** Left: Group average power spectral density plot (PSD, Mean ± SEM) in decibel (dB) of 5-HT ΔF/F signal during stimulation (red for NAc and blue for DS) versus non-stimulation epochs (grey). Right: 2 Hz signal-to-noise ratio quantification (SNR) of biosensor (470) and isosbestic (405) signal during stimulation versus non stimulation period (NAc : Two-way repeated measures ANOVA: Channel effect, F(_1,22_) = 0.013, P>0.05; Stimulation effect, F(_1,22_) = 8.501, **P<0.01; Channel vs Stimulation, F(_1,22_) = 14.06, **P<0.01; Bonferroni post hoc analysis, \*\*\**P*<0.001; DS: Two-way repeated measures ANOVA: Channel effect, F(_1,22_) = 16.70, ***P<0.001; Stimulation effect, F(_1,22_) = 12.98, **P<0.01; Channel vs Stimulation, F(_1,22_) = 19.34, ***P<0.01; Bonferroni post hoc analysis, \*\*\**P*<0.001). **(G)** Schematic of viral strategies used in SERT-Cre animals (*n* = 10) to simultaneously record DA release during optogenetic manipulation of DRN serotonergic neurons. **(H)** z-score group data (Mean ± SEM) of NAc (red) and DS (Blue) DA release evoked by optogenetic stimulation of DRN serotonergic neurons. **(I)** Quantification of AUC during optogenetic stimulation (Two-sided Mann-Whitney test, *P>*0.05) **(J,K)** Left: Group average power spectral density plot (PSD, Mean ± SEM) in decibel (dB) of DA ΔF/F signal during stimulation (red for NAc and blue for DS) versus non-stimulation epochs (grey). Right: 2 Hz signal-to-noise ratio quantification (SNR) of biosensor (470) and isosbestic (405) signal during stimulation versus non stimulation period (NAc : Two-way repeated measures ANOVA: Channel effect, F(_1,18_) = 6.53, *P<0.05; Stimulation effect, F(_1,18_) = 7.758, *P<0.05; Channel vs Stimulation, F(_1,18_) = 12.07, **P<0.01; Bonferroni post hoc analysis, \*\*\**P*<0.001; DS: Two-way repeated measures ANOVA: Channel effect, F(_1,18_) = 60.76, ***P<0.001; Stimulation effect, F(_1,18_) = 15.27, ***P<0.001; Channel vs Stimulation, F(_1,18_) = 13.57, **P<0.01; Bonferroni post hoc analysis, \*\*\**P*<0.001).

DR stimulation produced large 5-HT increases in both striatal regions (Figure 3C,D). The response developed throughout the stimulation period and remained elevated after stimulation ended. Spectral analysis also revealed increased power at 2 Hz restricted to the biosensor channel in both the NAc and DS, showing that striatal 5-HT signals followed the temporal structure of DR stimulation (Figure 3E,F).

By contrast, DR stimulation produced no sustained increase in the mean DA signal in either region (Figure 3G–I). Nevertheless, the DA traces contained small oscillatory responses phase-locked to stimulation (Fig 3H). Accordingly, spectral analysis revealed increased DA signal power at 2 Hz restricted to the biosensor channel in both the NAc and DS, with a more pronounced effect in the DS (Figure 3J,K). The absence of a corresponding signal in the isosbestic channel argues against direct optical contamination of the biosensor signal. Moreover, the same optical stimulation protocol applied in the VTA did not generate stimulation-locked 5-HT signals, making a nonspecific stimulation-related effect unlikely. DR serotonergic-neuron activation therefore strongly drove striatal 5-HT release while imposing a temporally locked component on DA signaling without noticeable increase of overall levels.

Together, these experiments reveal a directional asymmetric interaction between the two systems. VTA DA-neuron activity did not recruit striatal 5-HT release, whereas DR serotonergic-neuron activity weakly entrained striatal DA signals without producing a sustained increase in DA concentration.

## Discussion

Our results reveal a directionally asymmetric interaction between striatal DA and 5-HT signaling. Optogenetic activation of VTA DA neurons robustly increased DA release in the NAc but did not alter 5-HT signals in either the NAc or DS. Conversely, activation of DR serotonergic neurons strongly increased striatal 5-HT and produced a small DA component phase-locked to the stimulation, without generating a sustained increase in DA amplitude. Thus, under these experimental conditions, the two systems operate largely in parallel, with a weak influence of DR serotonergic activity on striatal DA signaling.

The modest DA response evoked by DR stimulation contrasts with reports of larger increases in striatal DA measured by microdialysis (24). Methodological differences may account for this discrepancy. Microdialysis integrates transmitter levels over relatively long sampling intervals and therefore emphasizes sustained changes, whereas genetically encoded sensors resolve rapid fluctuations in neuromodulator release. Differences in stimulation parameters, targeted neuronal populations and recording sites may also contribute. Our approach provides high temporal resolution and molecular selectivity, but it may be less sensitive to small, slowly accumulating changes in extracellular DA. The two methods may therefore capture different components of serotonergic modulation of DA transmission rather than yield directly contradictory results (25–27).

Although DR stimulation did not produce a sustained DA increase, it reliably enhanced DA-signal power at the stimulation frequency, particularly in the DS. This phase-locked component indicates that DR activity can influence dopaminergic transmission on a rapid timescale. One possible mechanism involves DR serotonergic neurons that co-release glutamate and project to the VTA, where they could transiently recruit DA neurons and promote striatal DA release (28). Alternatively, DR stimulation may engage polysynaptic pathways or local striatal mechanisms that modulate DA terminals. The present experiments do not distinguish among these possibilities.

Our anatomical tracing further revealed a rostrocaudal organization of DR neurons projecting to the NAc, DS and VTA. This organization supports the presence of partially segregated serotonergic output channels. However, the tracing data do not establish that this topography explains the functional asymmetry observed with optogenetic stimulation. Addressing that possibility will require projection-specific manipulation of defined DR subpopulations.

Several limitations constrain the interpretation of our findings. First, the stimulation protocol strongly and synchronously recruited genetically defined neuronal populations and may not reproduce their endogenous firing patterns during behavior. Second, we measured transmitter dynamics in two striatal regions but did not test other targets in which DA–5-HT interactions may differ. Third, the absence of a detectable 5-HT response after VTA stimulation does not exclude receptor-mediated interactions, slower modulatory effects or changes below the sensitivity of the sensors. Fourth, formally, visual perception of the red laser light, may have indirectly evoked dopamine transients. Finally, our experiments establish the directionality of acute release coupling but do not address how this interaction changes during learning, drug exposure or disease.

Together, these findings refine models of striatal monoamine signaling. Rather than functioning as tightly coupled or uniformly opponent systems, DA and 5-HT appear to follow largely parallel release pathways in the striatum. DR serotonergic activity can nevertheless impose a weak, temporally precise influence on DA signaling, revealing limited and directionally asymmetric crosstalk between the two systems.

## Methods

### Key Resources Table

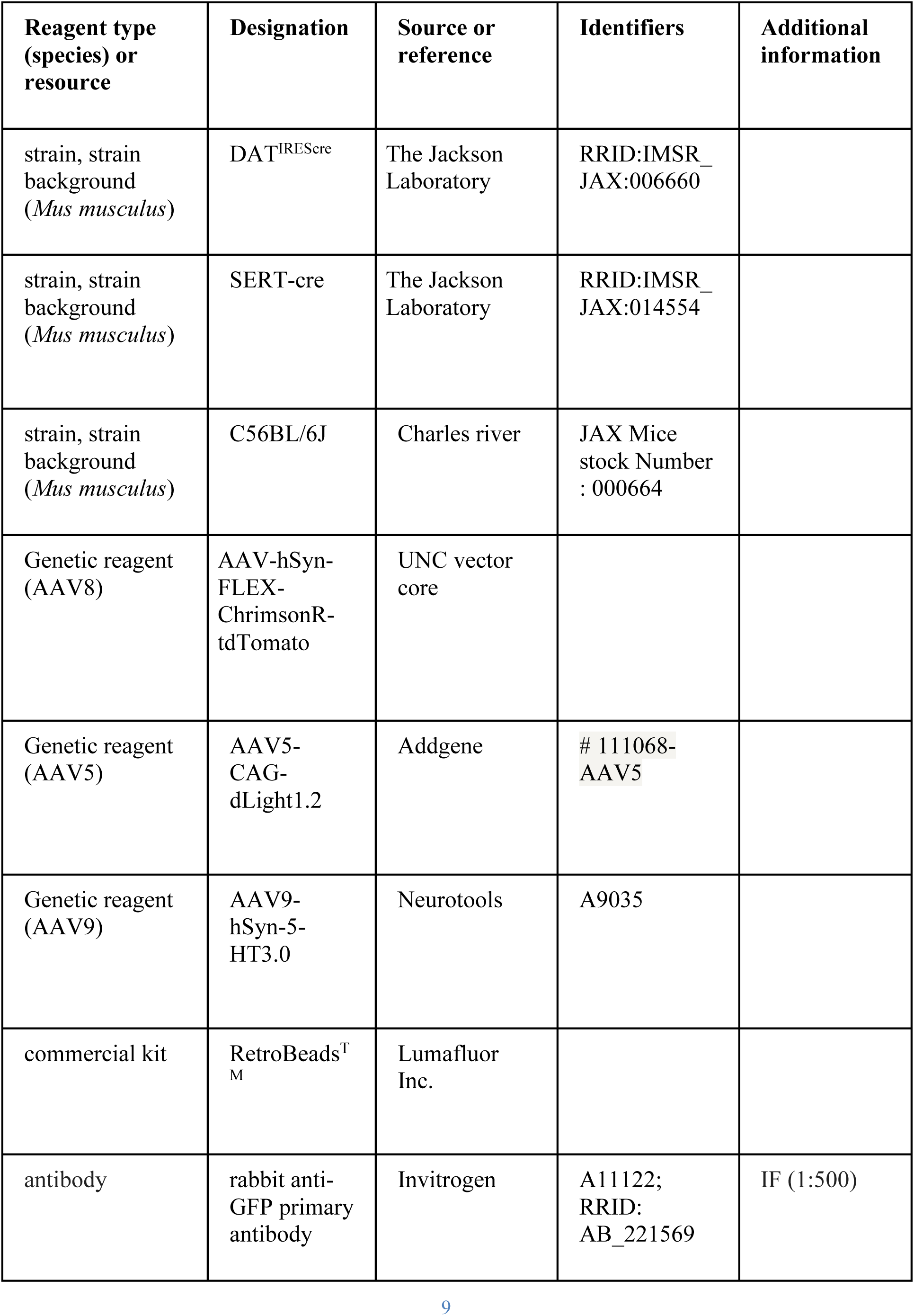
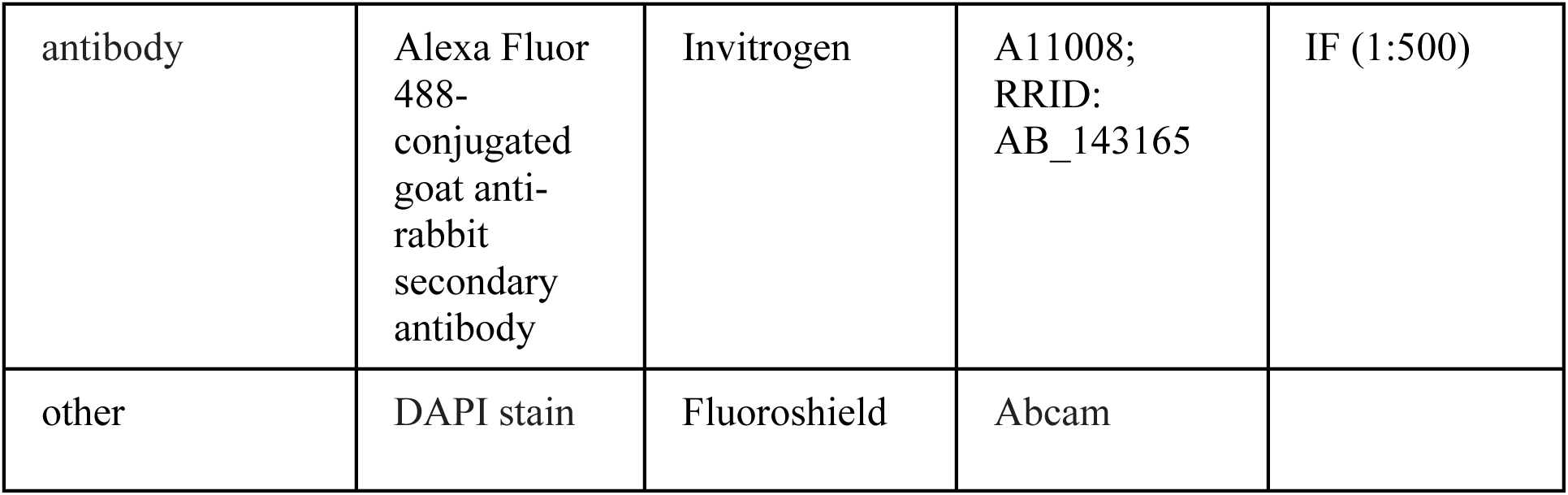

### Animals

DAT-IRES-Cre (B6.SJL-Slc6a3tm1.1(cre)Bkmn/J) and SERT-Cre (B6.129(Cg)-Slc6a4tm1(cre)Xz/J) mice were obtained from The Jackson Laboratory. C57BL/6J mice were purchased from Charles River Laboratories. Mice were allowed to habituate to the animal facility for 7 days after arrival. Male and female mice aged 8–12 weeks were group-housed in a temperature- and humidity-controlled environment under a 12-h light/dark cycle, with food and water available ad libitum. Body weights and sexes were balanced across groups whenever possible. All behavioral procedures were performed during the light phase. All procedures were approved by the Institutional Animal Care and Use Committee of the University of Geneva and the Animal Welfare Committee of the Canton of Geneva, in accordance with Swiss law (GE131A_33975).

### Viral injections and optical-fiber implantation

Mice aged 8–12 weeks were deeply anesthetized with isoflurane (Attane; 3% for induction and 1.5% for maintenance, in O₂) delivered using a compact anesthesia station (Minerve) and secured in a stereotaxic frame (Stoelting). Body temperature was maintained at 37 °C using a temperature control system, and Lacryvisc (Alcon, Switzerland) was applied to prevent corneal drying. Lidocaine was applied to the scalp before incision. The skull was exposed to visualize bregma and lambda, and a small craniotomy was made above each target region. Viruses and retrobeads were delivered through glass micropipettes.

To express the excitatory opsin, AAV8-hSyn-FLEX-ChrimsonR-tdTomato (UNC Vector Core) was injected into the ventral tegmental area (VTA; anteroposterior [AP], −3.28 mm; mediolateral [ML], −0.90 mm; dorsoventral [DV], −4.30 mm; 10° lateral angle) of DAT-Cre mice or into the dorsal raphe (DR; AP, −4.50 mm; ML, +0.55 mm; DV, −3.30 mm; 10° lateral angle) of SERT-Cre mice. To record dopamine (DA) or serotonin (5-HT) release, AAV5-CAG-dLight1.2 (Addgene, 111068) or AAV9-hSyn-5-HT3.0 (Neurotools, A9035), respectively, was injected into the nucleus accumbens (NAc; AP, +1.60 mm; ML, ±1.50 mm; DV, −4.37 mm; 10° lateral angle) or dorsal striatum (DS; AP, +0.80 mm; ML, ±1.65 mm; DV, −3.30 mm).

For anatomical tracing experiments, green retrobeads were injected into the VTA together with red retrobeads injected into either the NAc or DS (Lumafluor). The injection volume was 350 nL for viral vectors and 150 nL for retrobeads. After each injection, the pipette was left in place for 10 min to allow diffusion and minimize backflow.

During the same surgical procedure, three screws were secured to the skull to stabilize the implants. For DA and 5-HT recordings, optical fibres (400-µm diameter; MFC_400/430_0.48_4mm_ZF2.5(G)_FLT; Doric Lenses) were implanted 150 µm above the viral injection sites in the NAc and DS. A second optical fibre (200-µm diameter; FOC-W-1.25-200-0.37-5.0; Inper) was implanted 300 µm above the viral injection site in the VTA or DR for optogenetic stimulation. The optical fibres were secured to the skull with dental cement. The skin was sutured and disinfected, after which mice received an intraperitoneal injection of 500 µL saline. Paracetamol (2 mg/mL in the drinking water) was provided for 2–4 days after surgery. Mice were allowed to recover for 7 days before handling began.

### Behavioral paradigm

Following 4–5 weeks of viral expression, mice were habituated to handling, connection cables, and intraperitoneal saline injections on three consecutive days to minimize stress associated with the experimental procedures. On the day of the experiment, mice were connected to the optogenetic-stimulation cable and fibre-photometry patch cords and placed in a custom-built transparent open-field arena (40 × 40 × 30 cm). They were allowed to habituate to the arena for 3 min before recording began. For cocaine experiments, a 5-min baseline was recorded, followed by an intraperitoneal injection of cocaine (15 mg/kg) and 40 min of post-injection recording. For optogenetic-stimulation experiments, a 5-min baseline was recorded, after which mice received 20 stimulation trials of 30 bursts. Trials were delivered using randomized intertrial intervals of 20–40 s for DA recordings and 120–180 s for 5-HT recordings.

### Optogenetic stimulation

Mice were connected via patch cords (MFO-F-W1.25-200-0.37-100; Inper) to a rotary joint (FRJ_1×2_FC-2FC; Doric Lenses, Quebec, Canada) suspended above the open-field arena. A second patch cord connected the rotary joint to an orange diode-pumped solid-state laser (SDL-593-100 mW; Shanghai Dream Lasers, Shanghai, China) positioned outside the arena. Laser power, measured at the end of each patch cord, was typically 15–20 mW. Laser output was controlled by a mechanical shutter (SR474 driver with SR476 shutter head; Stanford Research Systems) aligned using a connectorized mechanical-shutter adaptor (Doric Lenses). Each stimulation trial consisted of 30 bursts delivered at 2 Hz over 15 s. Each burst comprised five 10-ms light pulses delivered at 20 Hz over 250 ms, with bursts repeated every 500 ms.

### Fibre-photometry recordings and analysis

Dual-fibre photometry was performed as previously described (29). During recordings, excitation light (470 nm; M470F3, Thorlabs) and isosbestic control light (405 nm; M405FP1, Thorlabs) were passed through excitation filters and focused onto a patch cord. For NAc recordings, the 470- and 405-nm LEDs were sinusoidally modulated at 271 and 331 Hz, respectively; for DS recordings, they were modulated at 425 and 375 Hz, respectively. Light was passed through a minicube (FMC4_AE(405)_E(450-490)_F(500-550)_S; Doric Lenses) and an optical-fibre patch cord (MFP_400/430/1100-0.48_4m_FC-ZF2.5; Doric Lenses) connected to the chronically implanted fibre (MFC_400/430-0.48_6mm_ZF2.5(G)_FLT; Doric Lenses). Light intensity at the tip of the patch cord was approximately 0.25 mW. Emitted light travelled back through the same fibres to the minicube and a photoreceiver (Newport 2151; Doric Lenses), after which the signal was digitized, demodulated, and stored using an RZ5P signal processor (Tucker-Davis Technologies). Signals were sampled at 101.725 Hz and low-pass filtered online at 20 Hz.

Data were analyzed using MATLAB R2024 (MathWorks). For each recording, the baseline signal obtained with 405-nm excitation was linearly regressed onto the signal obtained with 470-nm excitation and scaled to the 470-nm signal. ΔF/F was calculated as (F₄₇₀ − fitted F₄₀₅)/fitted F₄₀₅. The ΔF/F signal was then converted to a z score using either the 5-min pre-injection period for cocaine experiments or the 10-s pre-stimulation period for optogenetic experiments as the baseline. For optogenetic experiments, z scoring was performed separately for each trial to account for intertrial variability in fluorescence intensity. Normalized signals were averaged across trials for each mouse, and group data are presented as the mean across mice with the standard error of the mean (SEM).

The normalized ΔF/F signal and the raw 470- and 405-nm fluorescence signals were analyzed separately. Signals were first detrended and high-pass filtered using a fourth-order infinite impulse response filter with a cutoff frequency of 0.5 Hz to remove slow baseline fluctuations. Recordings were segmented into stimulation and non-stimulation epochs according to the onset and offset of each stimulation train. Power spectral density (PSD) was estimated for stimulation and non-stimulation epochs using Welch’s method with a 4-s window, 50% overlap, and a fast Fourier transform length equal to the next power of two. Analyses included frequencies from 0.5 to 10.5 Hz. The signal-to-noise ratio (SNR) at the stimulation frequency was calculated as the mean spectral power within the 1.5–2.5-Hz band divided by the mean power at the remaining frequencies from 0.5 to 10.5 Hz, excluding the stimulation band.

### Histological analysis and retrobead quantification

At the end of the experiments, mice were deeply anesthetized with pentobarbital (300 mg/kg, intraperitoneally) and transcardially perfused with 4% (w/v) paraformaldehyde (PFA) in phosphate-buffered saline (PBS). Brains were post-fixed overnight at 4 °C and sectioned using a vibratome (VT1200; Leica). For immunohistochemistry, 50-µm-thick sections were washed three times in PBS for 5 min and blocked for 1 h at room temperature in PBS containing 10% bovine serum albumin and 0.5% Triton X-100. Sections were incubated overnight at 4 °C with rabbit anti-GFP primary antibody (1:500; Invitrogen A11122; RRID: AB_221569) diluted in blocking solution. This antibody was used to amplify the GFP-derived fluorescence of the genetically encoded sensors and thereby verify sensor expression and fibre placement. Sections were then washed four times in PBS for 15 min and incubated for 2 h at room temperature with Alexa Fluor 488-conjugated goat anti-rabbit secondary antibody (1:500; Invitrogen A11008; RRID: AB_143165) diluted in blocking solution. After four additional 15-min washes in PBS, sections were mounted using DAPI-containing mounting medium (Fluoroshield; Abcam). For retrobead experiments, 60-µm-thick sections were mounted directly using DAPI-containing mounting medium.

Sections containing the VTA, NAc, DS, and DR were imaged using a Zeiss Axioscan Z1 equipped with a Plan-Apochromat 10×/0.45 numerical-aperture objective and filter sets for DAPI (445/50-nm emission), eGFP (525/50-nm emission), and cyanine 3 (Cy3; 605/70-nm emission). Cells were automatically quantified in QuPath using the cell-detection plugin.

### Statistical analysis

For each experiment, a minimum of two replications were conducted by experimenters. Statistical analyses were performed using GraphPad Prism 11. The significance threshold was set at α = 0.05. Normality was assessed using the D’Agostino–Pearson test. Mann–Whitney or Friedman tests were used for non-Gaussian data, as appropriate. Multiple comparisons were analyzed using mixed-factor analyses of variance (ANOVAs) with the appropriate between- and within-subject factors (e.g. biosensor/isosbestic channels). When significant main effects or interactions were detected (P < 0.05), post hoc comparisons were performed using two-tailed Student’s t tests with Bonferroni correction, as appropriate.

## Acknowledgments

We thank all the Lüscher lab members for their comments on the manuscript. This work was supported by an advanced grant from the European Research Council (F-Addict) and DEEPER project (European community) to C.L.

## Author information

Conceptualization: FC, YL, CL Investigation: FC, YL, JET, AH Formal Analysis: FC, YL Visualization: FC, YL, CL Funding acquisition: CL Supervision: CL Writing – original draft: FC, YL, CL Writing – review & editing: FC, YL, CL

## Data availability statement

The datasets generated and analyzed for the current study are available in the Zenodo repository DOI : <u>10.5281/zenodo.21493873</u>. Source data and code are provided also within the repository.

